# Uncovering Cross-Cohort Molecular Features with Multi-Omics Integration Analysis

**DOI:** 10.1101/2022.11.10.515908

**Authors:** Min-Zhi Jiang, François Aguet, Kristin Ardlie, Jiawen Chen, Elaine Cornell, Dan Cruz, Peter Durda, Stacey B. Gabriel, Robert E. Gerszten, Xiuqing Guo, Craig W. Johnson, Silva Kasela, Leslie A. Lange, Tuuli Lappalainen, Yongmei Liu, Alex P. Reiner, Josh Smith, Tamar Sofer, Kent D. Taylor, Russell P. Tracy, David J. VanDenBerg, James G. Wilson, Stephen S. Rich, Jerome I. Rotter, Michael I. Love, Laura M. Raffield, Yun Li, NHLBI Trans-Omics for Precision Medicine (TOPMed) Consortium, TOPMed Analysis Working Group

**Affiliations:** Department of Applied Physical Sciences, University of North Carolina at Chapel Hill, Chapel Hill, NC, United States; Illumina Artificial Intelligence Laboratory, Illumina, Inc., San Diego, CA, United States; The Broad Institute of MIT and Harvard, Cambridge, MA, United States; Department of Biostatistics, University of North Carolina at Chapel Hill, Chapel Hill, NC, United States; Laboratory for Clinical Biochemistry Research, University of Vermont, Burlington, VT, United States; Department of Medicine, Cardiology, Beth Israel Deaconess Medical Center, Boston, MA, United States; Department of Pathology & Laboratory Medicine, University of Vermont, Colchester, VT, United States; Department of Medicine, Beth Israel Deaconess Medical Center, Boston, MA, United States; Department of Pediatrics, The Institute for Translational Genomics and Population Sciences, The Lundquist Institute for Biomedical Innovation at Harbor-UCLA Medical Center, University of California at Los Angeles, Torrance, CA, United States; Department of Biostatistics, University of Washington at Seattle, Seattle, WA, United States; New York Genome Center, New York, NY, United States; Department of Epidemiology, Department of Medicine, Division of Biomedical Informatics and Personalized Medicine, Lifecourse Epidemiology of Adiposity & Diabetes Center, Aurora, CO, United States; Department of Medicine, Cardiology and Neurology, Duke University Medical Center, Durham, NC, United States; Department of Epidemiology, University of Washington, Seattle, WA, United States; Northwest Genomic Center, University of Washington, Seattle, WA, United States; Department of Biostatistics, Harvard Medical School, Medicine-Brigham and Women’s Hospital, Boston, MA, United States; Department of Preventive Medicine, University of Southern California, Los Angeles, CA, United States; Center for Public Health Genomics, Department of Public Health Sciences, University of Virginia, Charlottesville, VA, United States; Department of Pediatrics, Genomic Outcomes, The Institute for Translational Genomics and Population Sciences, The Lundquist Institute for Biomedical Innovation at Harbor-UCLA Medical Center, University of California at Los Angeles, Torrance, CA, United States; Department of Genetics, Department of Biostatistics, University of North Carolina at Chapel Hill, Chapel Hill, NC, United States; Department of Genetics, University of North Carolina at Chapel Hill, Chapel Hill, NC, United States

## Abstract

Integrative approaches that simultaneously model multi-omics data have gained increasing popularity because they provide holistic system biology views of multiple or all components in a biological system of interest. Canonical correlation analysis (CCA) is a correlation-based integrative method. It was initially designed to extract latent features shared between two assays by finding the linear combinations of features – referred to as canonical vectors (CVs) – within each assay that achieve maximal across-assay correlation. Sparse multiple CCA (SMCCA), a widely-used derivative of CCA, allows more than two assays but can result in non-orthogonal CVs when applied to high-dimensional data. Here, we incorporated a variation of the Gram-Schmidt (GS) algorithm with SMCCA to improve orthogonality among CVs. Applying our SMCCA-GS method to proteomics and methylomics data from the Multi-Ethnic Study of Atherosclerosis (MESA) and Jackson Heart Study (JHS), we identified strong associations between blood cell counts and protein abundance. This finding suggests that adjustment of blood cell composition should be considered in protein-based association studies. Importantly, CVs obtained from two independent cohorts demonstrate transferability across the cohorts. For example, proteomic CVs learned from JHS explain similar amounts of blood cell count phenotypic variance in MESA, explaining 39.0% ~ 50.0% variation in JHS and 38.9% ~ 49.1% in MESA, similar transferability was observed for other omics-CV-trait pairs. This suggests that biologically meaningful and cohort-agnostic variation is captured by CVs. We further developed Sparse Supervised Multiple CCA (SSMCCA) to allow supervised integration analysis for more than two assays. We anticipate that applying our SMCCA-GS and SSMCCA on various cohorts would help identify cohort-agnostic biologically meaningful relationships between multi-omics data and phenotypic traits.

**Author Summary:** Comprehensive understanding of human complex traits may benefit from incorporation of molecular features from multiple biological layers such as genome, epigenome, transcriptome, proteome, and metabolome. CCA is a correlation-based method for multi-omics data which reduces the dimension of each omic assay to several orthogonal components – commonly referred to as canonical vectors (CVs). The widely-used SMCCA method allows effective dimension reduction and integration of multi-omics data, but suffers from potentially highly correlated CVs when applied to high-dimensional omics data. Here, we improve the statistical independence among the CVs by adopting a variation of the GS algorithm. We applied our SMCCA-GS method to proteomic and methylomic data from two cohort studies, MESA and JHS. Our results reveal a pronounced effect of blood cell counts on protein abundance, strongly suggesting blood cell composition adjustment in protein-based association studies may be necessary. Finally, we present SSMCCA which allows supervised CCA analysis for the association between one phenotype of interest and more than two assays. We anticipate that SMCCA-GS would help reveal meaningful system-level factors from biological processes involving features from multiple assays; and SSMCCA would further empower interrogation of these factors for phenotypic traits related to health and diseases.

## Introduction

In recent years, there has been rapid growth in high-dimensional multi-omics datasets (including DNA methylation, RNA-sequencing, metabolomics, proteomics, genomics, microbiome, etc). However, careful analyses with integrative methods are needed to fully utilize these rich datasets and provide mechanistic insights into health and disease related outcomes. While many methods have been published [1–3], few studies have evaluated these methods on large-scale datasets from human samples. In addition, despite quite a few successful examples of integrating two omics data-types [4–8], particularly detection of quantitative trait loci using genomic data, there are much fewer such examples of integrative analyses across more than two omics data types.

One promising method for using multi-omics data to explain phenotypic variation in health outcomes is canonical correlation analysis (CCA) [9]. CCA is a statistical technique to identify associations among two assays where each assay contains multiple variables. Specifically, CCA finds a linear combination of variables in each assay that leads to the maximal correlation of the two linear combinations. Principal component analysis (PCA) can be considered as a special case of CCA as the optimization objective is the same in the case that the same data is used for the two assays. CCA is a commonly adopted dimension reduction and information extraction method in genomic studies [1,10–13] as increasingly more modern genomic studies collect data from multiple assays.

An extension of CCA by Witten & Tibshirani [1] called sparse multiple CCA (SMCCA) allows for the input of multiple assays. We hypothesized that this method would be helpful for high-dimensional multi-omics data exploration and for understanding and extracting omics signatures that reflect biologically relevant variations. Specifically, we here leverage our CCA-based method extended from Witten & Tibshirani’s SMCCA to extract low-dimensional latent variables from high-dimensional multi-omics data and use them to explain phenotypic traits, focusing on blood cell indices, along with basic demographic and anthropometric characteristics. We perform CCA-based analyses in two studies with rich multi-omics data in hundreds of individuals, the Multi-Ethnic Study of Atherosclerosis (MESA) and the Jackson Heart Study (JHS).

## Results

### CCA Pipline

A typical CCA-based method generates orthogonal canonical vectors (CVs), which are low-dimensional summaries to represent latent variables underlying the multi-assay input data. **Fig 1** is a cartoon illustration where we have three assays (X, V, and Z) for three samples. For presentation brevity, we only show how we obtain the top 4 CVs. For each assay, CCA infers 4 vectors of weights (e.g., *W*_*X*1_, *W*_*X*2_, *W*_*X*3_, and *W*_*X*4_ for assay *X*), which leads to four CVs. For example, *CV*_*X*1_, the top CV for assay *X*, is obtained by *X* × *W*_*X*1_. The weights are inferred by maximizing the correlation of CVs across three assays. Note that in the rightmost CV matrices, each column of a CV matrix is one CV of the corresponding assay. In addition, CVs corresponding to the same column cross assays are expected to have maximal correlation (for instance, *CV*_*X*1_, *CV*_*Y*1_, *CV*_*Z*1_ are most correlated), while CVs in different columns are expected to be orthogonal or independent from each other in the same assay.

**Fig 1.**
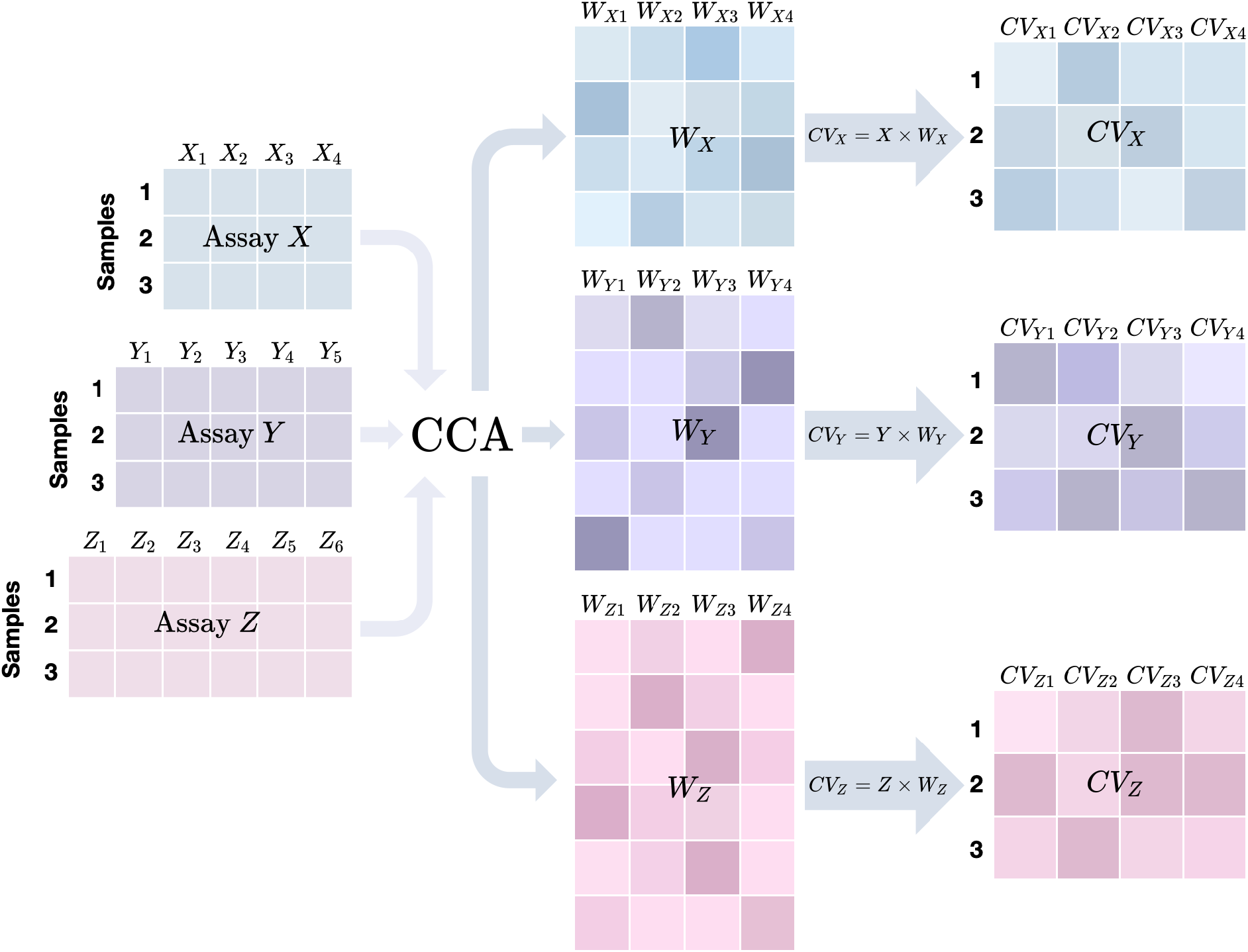
Cartoon illustration of a typical CCA-based method for three assays. X, Y, and Z are three assays with 4, 5, and 6 features respectively. When applying a CCA-based method on them to compute 4 **canonical vectors** (CVs), we would first get their **weight matrices** Wx, Wy, Wz, each of which contains 4 weight vectors. By multiplying each assay matrix (left panel) and its corresponding weight matrix (middle panel), we obtain the CV matrix for the assay (right panel) where each column corresponds to one CV.

### Leveraging CCA to identify and correct sample swaps

We first explored CCA for quality control (QC) purposes in multi-omics data. CCA, a multivariate statistical method, extracts correlated latent variations across multiple assays and thus naturally lends itself to multi-omics QC. We developed a CCA-based sample matching method (**Fig 2A-C**). For brevity, assume we have two assays *X* and *Y* with sample sizes *N_X_* and *N_Y_* (with *N* overlapping). First, using the *N* overlapping samples, we obtain top *K* (*K*=30 by default) CVs. Applying the learned CVs to all samples, we define similarity scores S(*X_i_, Y_i_*) = corr(*CV*_*X,i*_, *CV*_*Y,j*_) where *i* = 1,2,…, *N_X_*, and *CV*_*X*,i_ and *CV_Y,i_* are vectors of dimension *K*. We then construct a similarity score matrix, where the *i, j* entry is *S*(*X_i_, Y_j_*). Outliers are detected according to this similarity score matrix [14]. Supposing that a specific sample indexed by *p* is identified as an outlier, its similarity score, *S*(*X_p_, Y_p_*), would be significantly smaller than *S*(*X_m_, Y_m_*) where *m* indexes a “non-problematic” sample. We search the rest of the samples to identify a better mutual match (e.g., finding a sample *q* with S(*X_p_, Y_q_*) and S(*X_q_, X_p_*) comparable to S(*X_m_, Y_m_*)). We have found such an approach effective in identifying samples with mismatching ‘omic features and finding potential matches from other samples. **Fig 2D-F** shows an example from MESA where we swapped labels of samples # 82 and # 593 in their proteomics data (labels assigned at random, **Fig 2E**), while keeping their methylomics data labels intact. Similarly, **Fig 2F** shows the impact of swapping labels for these samples in the methylomics data (while keeping the protein data labels intact). The top CV alone can identify both as outliers when labels in either assay are swapped.

**Fig 2.**
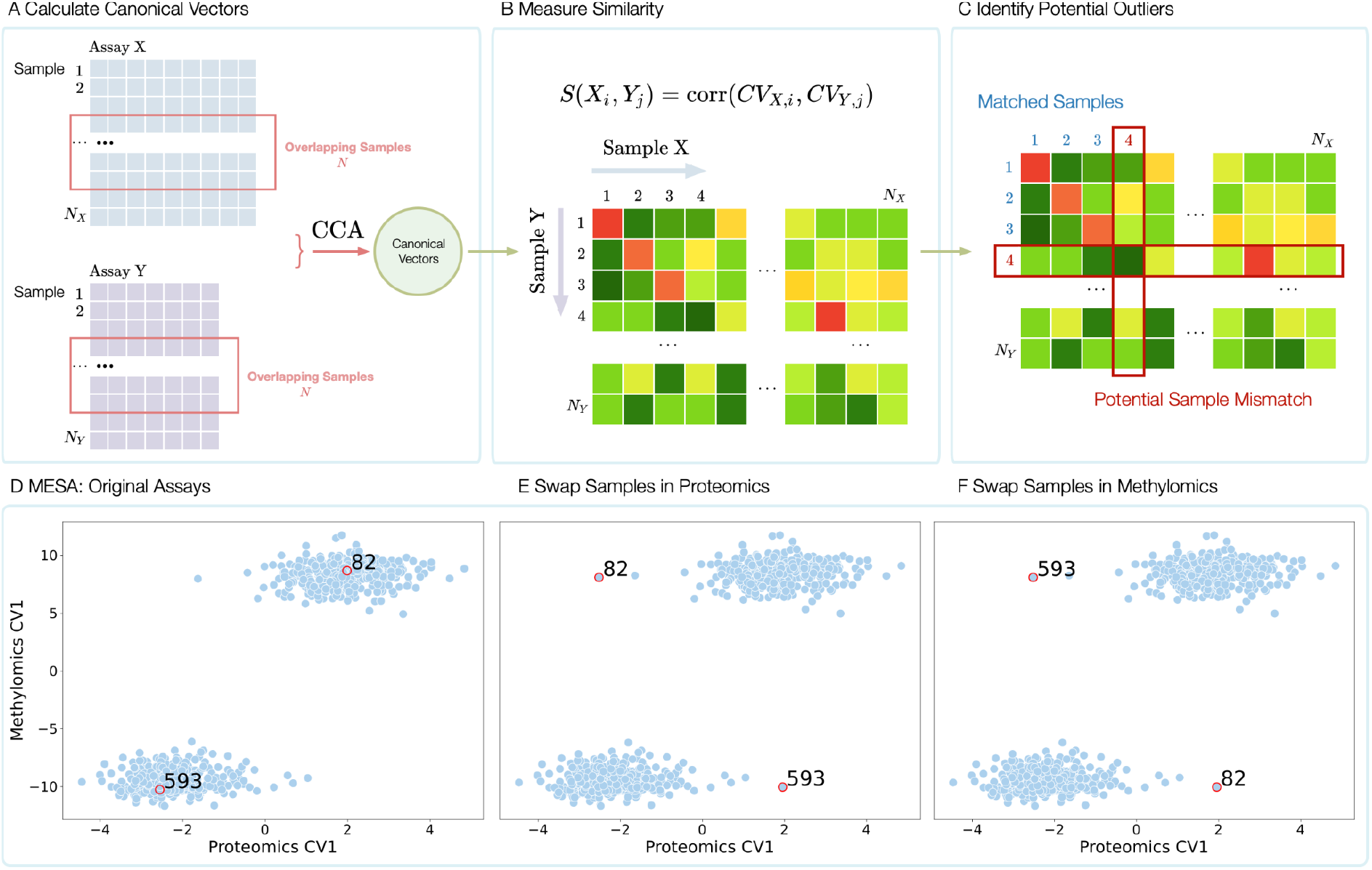
CCA-based method to identify and correct sample swaps. (**A-C**) Schematic of the method; (**D-F**) Proof-of-concept example in MESA. X-axis: top CV or CV1 from proteomics data for each sample; Y-axis: top CV or CV1 from methylomics data for each sample. The two clusters of points reflect the two sexes. We swap the labels for two samples (with example individual IDs 82 and 593), either their proteomics data (**Fig 2E**) or their methylomics data (**Fig 2F**).

### Modified Gram-Schmidt Algorithm Improves Orthogonality

SMCCA implemented in the PMA R package does not always provide the expected orthogonal CVs. For example, **Fig 3A-B** shows results from PMA’s implementation of unsupervised SMCCA when applied to MESA proteomics and methylomics data (detailed in Methods) where we observe extensive correlation among the CVs. In the presence of undesired correlated CVs, users will have to perform a secondary filtering step to generate a list of non-redundant CVs, or else variation in omics data captured by the later CVs may overlap with variance captured by former CVs. Therefore, we sought to improve orthogonality among generated CVs for capturing distinct information from the integrated multi-omics data. Specifically, we follow the Gram–Schmidt (GS) strategy [15] which generates CVs sequentially by progressively subtracting the previous CV from the input matrices (detailed in Methods). **Fig 3C-D** shows substantially improved orthogonality among the CVs when applied to the same MESA proteomics and methylomics data.

**Fig 3:**
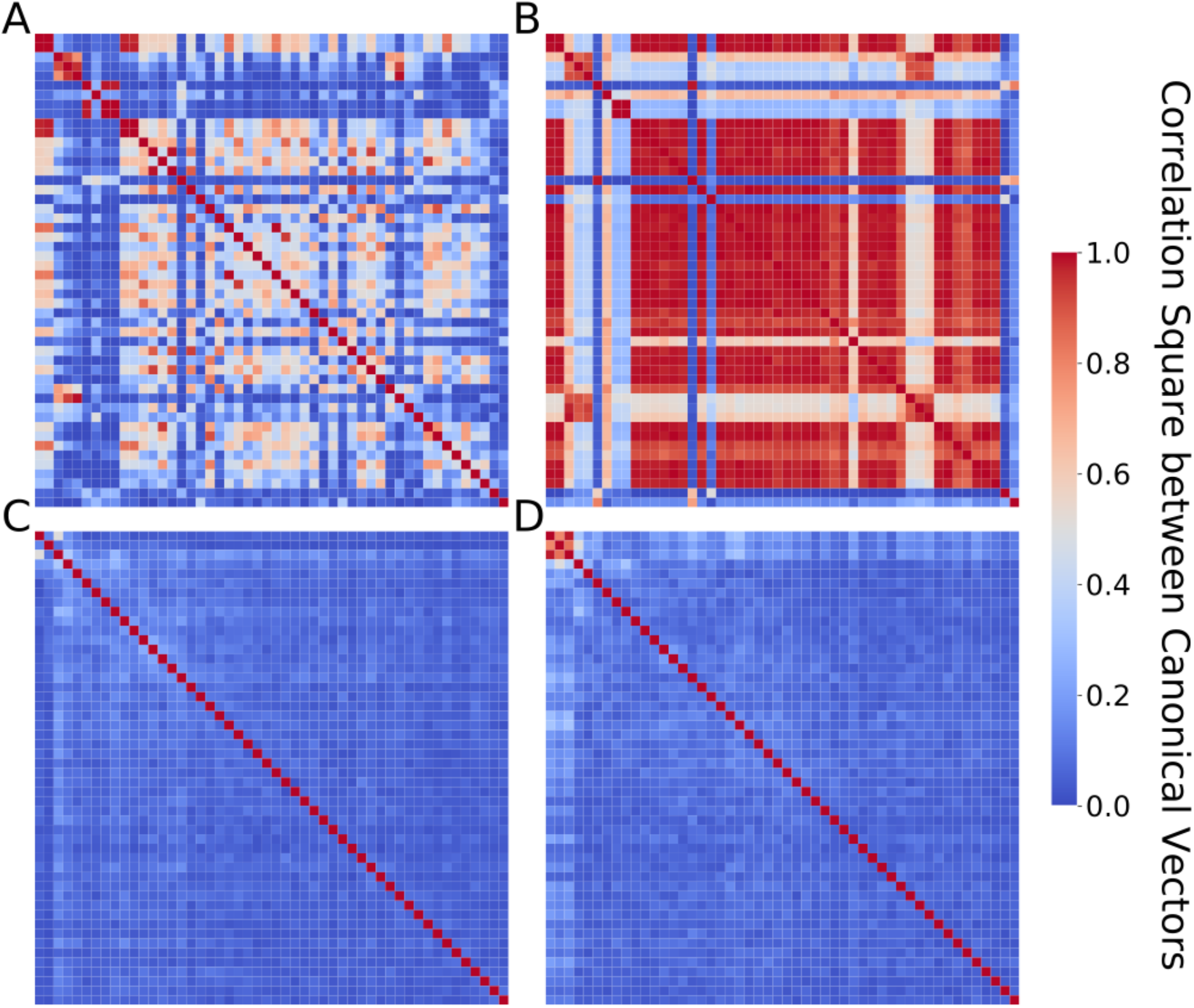
Improved orthogonality among CVs by adopting the Gram–Schmidt (GS) strategy. CVs are inferred from MESA proteomics and methylomics data using unsupervised SMCCA. Each row and column represent one CV, ranging from CV1 to CV50. **(A-B)** Results from PMA implementation. **A** shows CVs from proteomics data, and **B** from methylomics data. **(C-D)** Results from our SMCCA-GS implementation. **C** from proteomics data, and **D** from methylomics data.

### Proteomics CVs explain considerable amounts of variation in blood cell counts

We also applied our implementation to proteomics and methylomics data in JHS. As these unsupervised CVs are anticipated to capture shared latent variables underlying the proteomics and methylation datasets, we hypothesized that the CVs may explain a non-negligible amount of variation in various phenotypes. Our primary phenotypes of interest in this work are blood cell traits, including white blood cell count (WBC), red blood cell count (RBC) and platelet count (PLT). We also considered age and body mass index (BMI), as “control” phenotypes which have been widely reported to explain considerable variability in proteomics and methylomics data. For each of the five outcome phenotypes, we fit regression models to estimate the percent of variation explained by the top 50 CVs from each of the two omics data, namely proteomics and methylomics (detailed in Methods). For each cohort (MESA or JHS), we had two sets of CVs, one derived from the cohort’s own omics data, the other derived from applying the CV weights inferred from the other cohort.

We found that top CVs, from each of the two omics data, explain considerable amounts of variation in almost all of the outcomes evaluated (**Fig 4**). For example, top 50 methylomics CVs inferred in JHS explained 0.72, 0.35, 0.37, 0.34, 0.30 variation in age, BMI, WBC, RBC, and PLT respectively, in JHS (**Fig 4A**). We also observe high transferability between MESA and JHS. For example, the top 50 methylomics CVs inferred in MESA explained similar amounts of variation in RBC: 0.33 in MESA (itself) (**Fig 4C**) and 0.30 when applied to JHS (**Fig 4D**). Such high transferability suggests that latent variables learned by CCA might reflect essential biological processes shared across cohorts. We also note that these *r*^2^’s from methylomics data were most likely under-estimated because the CVs were constructed using the top 10,000 most variable CpG sites (see Methods) instead of the entire over 800,000 sites, for computational reasons. These findings are not surprising: for instance, blood cell composition (notably for white blood cell subtypes) has been long known to influence the methylome. For that reason, in epigenome-wide association studies (EWAS), it has been standard practice to first estimate the leukocyte proportions from methylomics data and adjust for these cell type proportions in subsequent association analysis[16]. Given shared precursors for all hematological cell types, we found it relatively unsurprising that RBC and PLT also had a high percent variation explained by methylomics CVs. Similarly, age [17] and BMI [18] have been known to explain substantial variability in methylomics data, and are commonly adjusted for as covariates.

**Fig 4.**
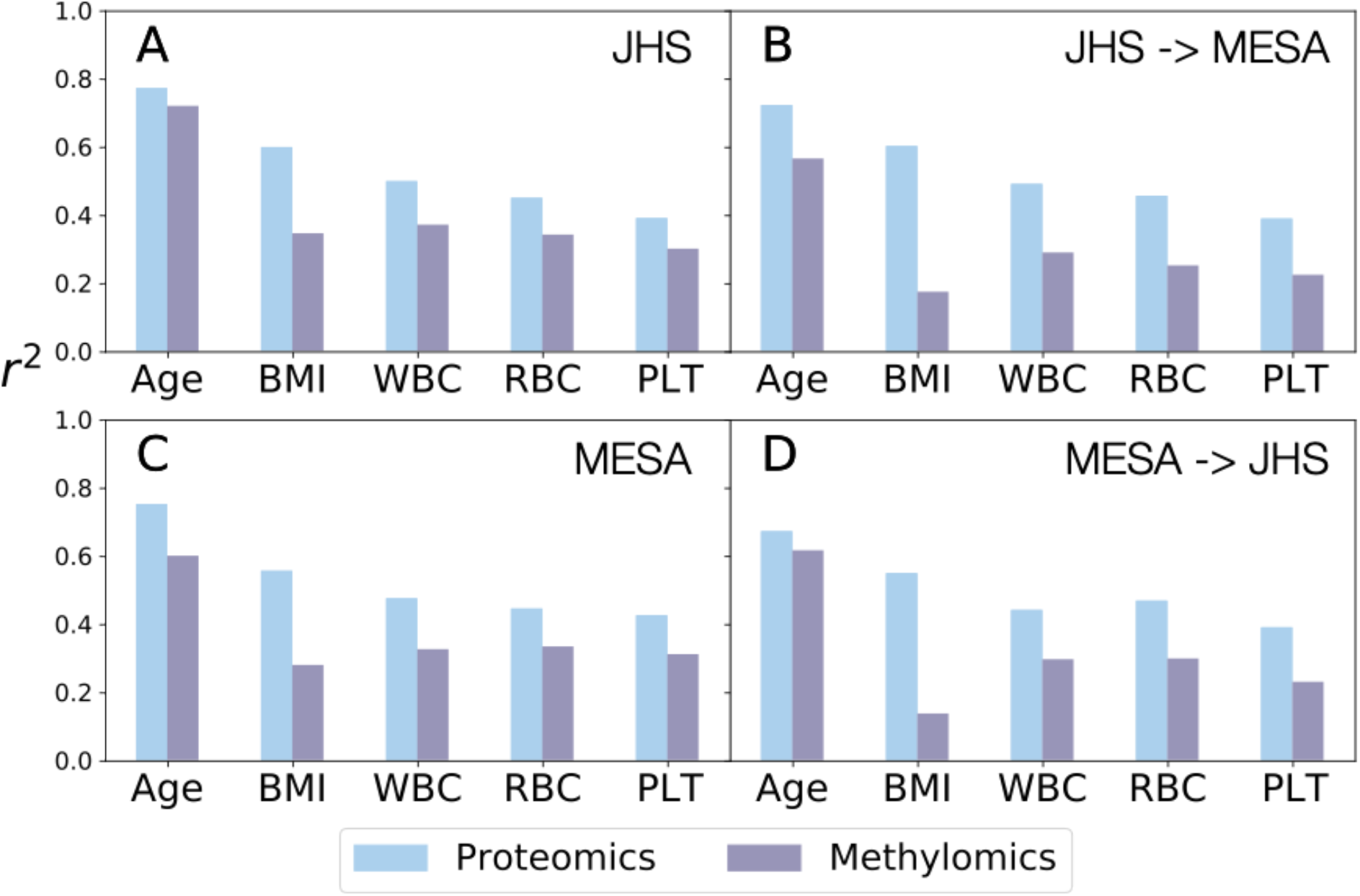
Proportion of variation in outcomes explained by CVs. **(A)** CVs were inferred using proteomics and methylomics in JHS. The top 50 CVs were used to calculate the *r*^2^ (Y-axis) for each outcome (X-axis). **(B)** We obtained CVs in JHS by applying the weights inferred from MESA, and then calculated *r*^2^ in the same way as in A. **(C)** CVs were inferred using proteomics and methylomics in MESA. **(D)** CVs were obtained in MESA by applying the weights inferred from JHS.

More interestingly, the amounts of variation in various outcomes explained by top 50 *proteomics* CVs are even higher, ranging 0.39-0.77 in JHS and 0.39-0.72 in MESA. Large *r*^2^ for age and BMI are expected since both have been reported to rather broadly affect protein profiles [19,20]. Strikingly, *r_2_* for blood cell traits are also considerable, and comparable to BMI, 0.50, 0.45, 0.39 respectively for WBC, RBC and PLT in JHS using CVs inferred in JHS. Confirmatorily, when applying CV weights inferred from MESA to JHS, we obtained similar *r*^2^’s: 0.44, 0.47, 0.39 for WBC, RBC and PLT respectively. Similar patterns were also observed in MESA. These considerable amounts of variations in blood cell counts explained by top proteomics CVs have important implications for association studies involving proteomics data: we should consider adjusting for blood cell proportions in these association studies, under the same rationale in EWAS (variability driven by blood cell subtype abundance is likely not of interest for many disease outcomes of interest whose association with proteomics data is being examined).

### CVs vs Principal Components (PCs)

Although CVs are inferred *jointly* from multi-omics data, we have focused on analyzing CVs from each omics data type *separately* for their predictive power of outcomes of interest. Thus, we naturally are interested in comparing the CCA-based approach with the standard PCA approach since we can obtain PCs separately for each omics data. Note first that we expect larger and more assay-specific batch effects in JHS than MESA. For example, JHS proteomics data was generated in 3 batches [21], and separately from the methylomics data. In contrast, MESA proteomics and methylomics data were all generated through the MESA TOPMed pilot over a short time period [22,23]. Results shown in **Fig 5** supported our expectations: overall we observe that a lower number of JHS inferred CVs are needed to explain the outcomes with higher r2 compared to JHS inferred PCs, indicating that top CVs inferred from JHS data tend to capture biological variations while top PCs tend to reflect more assay-specific technical variations. The contrasts are most pronounced with age and WBC for proteomics data, and with age for methyomics data. For example, in JHS, proteomics-CV1 explained 33% variation in WBC (blue + on the leftmost side of **Fig 5A3**) while proteomics-PC1 only explained 7.7% (purple + on the leftmost side of **Fig 5A3**). This noticeable advantage continued until ~20 CVs/PCs. For instance, the top 15 proteomics-CVs in JHS explained 44% variation in WBC (blue X in **Fig 5A3**) while top 15 proteomics-PCs only explained 29% (purple X in **Fig 5A3**). Similar advantages of CVs over PCs were observed in MESA, but were less pronounced as expected due to the smaller and less assay-specific batch effects in MESA. Reassuringly, applying JHS inferred CV weights to MESA showed advantages similar to those in JHS, more pronounced than using CVs inferred in MESA itself, further demonstrating the power of CVs to capture biologically relevant variations under the presence of assay-specific batch effects.

**Fig 5.**
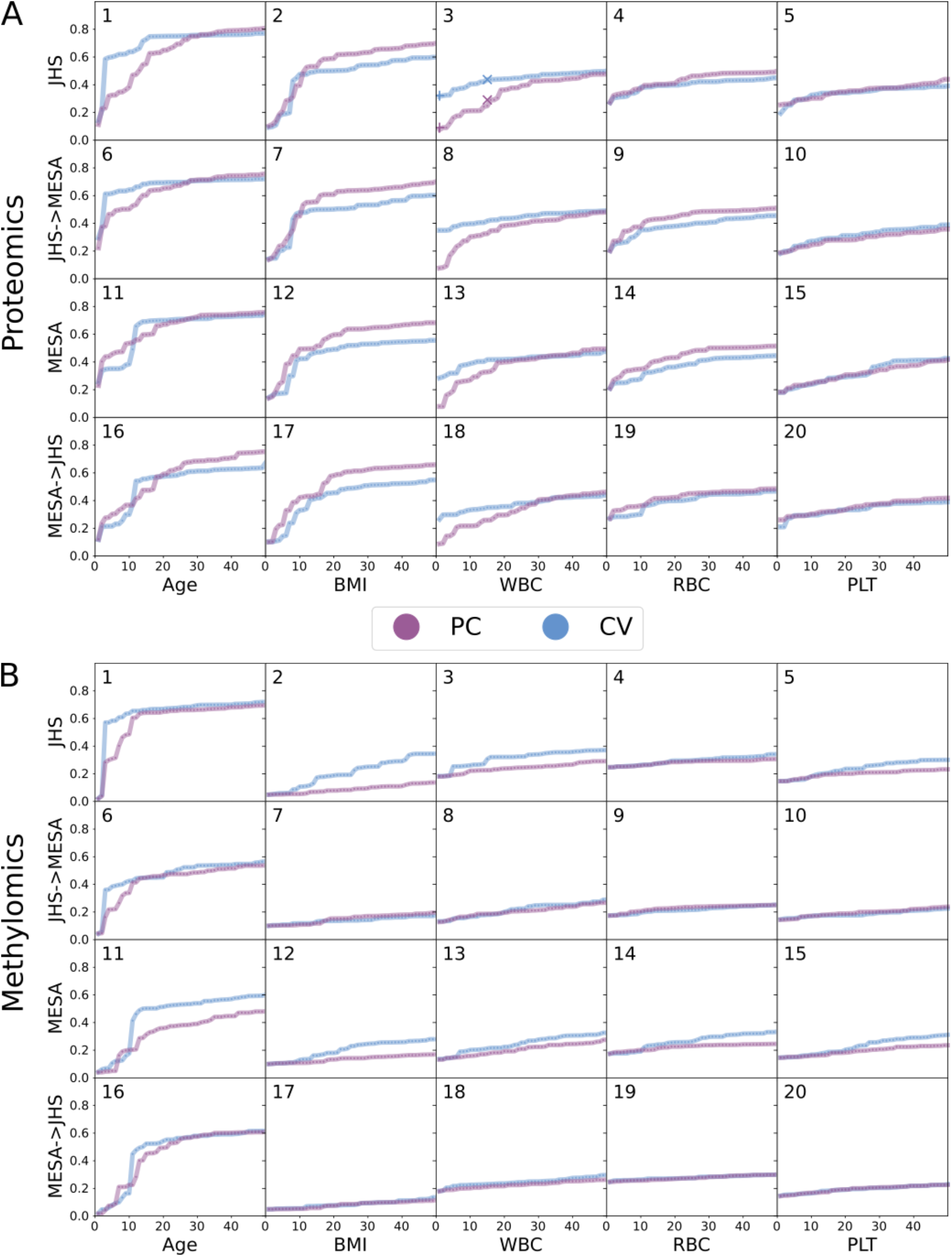
Comparison of *r*^2^, PCs vs CVs. Each column corresponds to one outcome. Within each panel, top row (JHS) shows results in JHS using JHS-inferred CVs. Second row (JHS->MESA) shows results in MESA, also using JHS-inferred weights. Third row (MESA) shows results in MESA, this time using MESA-inferred CVs. Last row (MESA->JHS) shows results in JHS, also using MESA-inferred weights. **(A)** Proteomics. **(B)** Methylomics. In each sub-figure, X-axis indicates the number of CVs or PCs used and Y-axis the proportion of variation explained in the outcome (i.e., *r*^2^).

### Supervised Sparse Multiple CCA

#### Extending Supervised Sparse CCA to Supervised Sparse Multiple CCA

So far we have generated and evaluated unsupervised CCA where the CVs are inferred from multi-omics data only, without considering any outcomes of interest. Although we assessed the relationship between unsupervised CVs and several outcomes of interest, the CVs themselves were inferred without knowledge of the outcomes. In practice, when we are primarily interested in a particular outcome, supervised approaches can be more effective and powerful. The PMA R package implements a sparse supervised CCA (SSCCA) method. However, this implementation only accepts two omics data at a time, which limits our capabilities in real datasets where there are more than two assays. For instance, in both MESA and JHS, we also have whole genome sequencing (WGS) data [24]. We implemented a sparse supervised multiple CCA (SSMCCA) method to accomodate more than two assays of omics data. Our implementation follows the idea in Witten et al., (2009) where a feature selection step is performed within each assay to retain (by default) top ~80% features most correlated with the outcome of interest. Features selected from each assay form new input matrices to which we then apply our implementation of unsupervised SMCCA with the adapted Gram-Schmidt algorithm.

To ensure our SSMCCA implementation generates sensible supervised CVs, we first compared results from PMA’s SSCCA implementation, when there are two assays of data. Specifically, we compared correlations between inferred supervised CV1 and the corresponding outcomes of interest. We compared SSCCA and our SSMCCA by running two methods with 100 different random seeds and for each seed, testing the variation of each outcome explained by supervised proteomics CVs and supervised methylomics CVs (**Fig 6**). We found that in most cases, the amount of variation in outcomes captured by SSMCCA CVs is comparable or significantly higher than SSCCA, indicated by large red circles. For example, **Fig 6A** third row third column, SSMCCA proteomics CV1 explains 4.17% variation in PLT in MESA, while SSCCA 3.48% (significance of their difference is p-value = 8 × 10^-9^). In a few cases, the amount of variation captured by SSMCCA CV1 is significantly smaller than SSCCA CV1. For example **Fig 6B** row 2 column 1, although the difference in terms of percent variation explained in RBC by SSCCA vs SSMCCA methylomics CV1 is highly significant (p-value = 3× 10^-28^), the absolute difference (4.72 × 10^-8^) is tiny.

**Fig 6.**
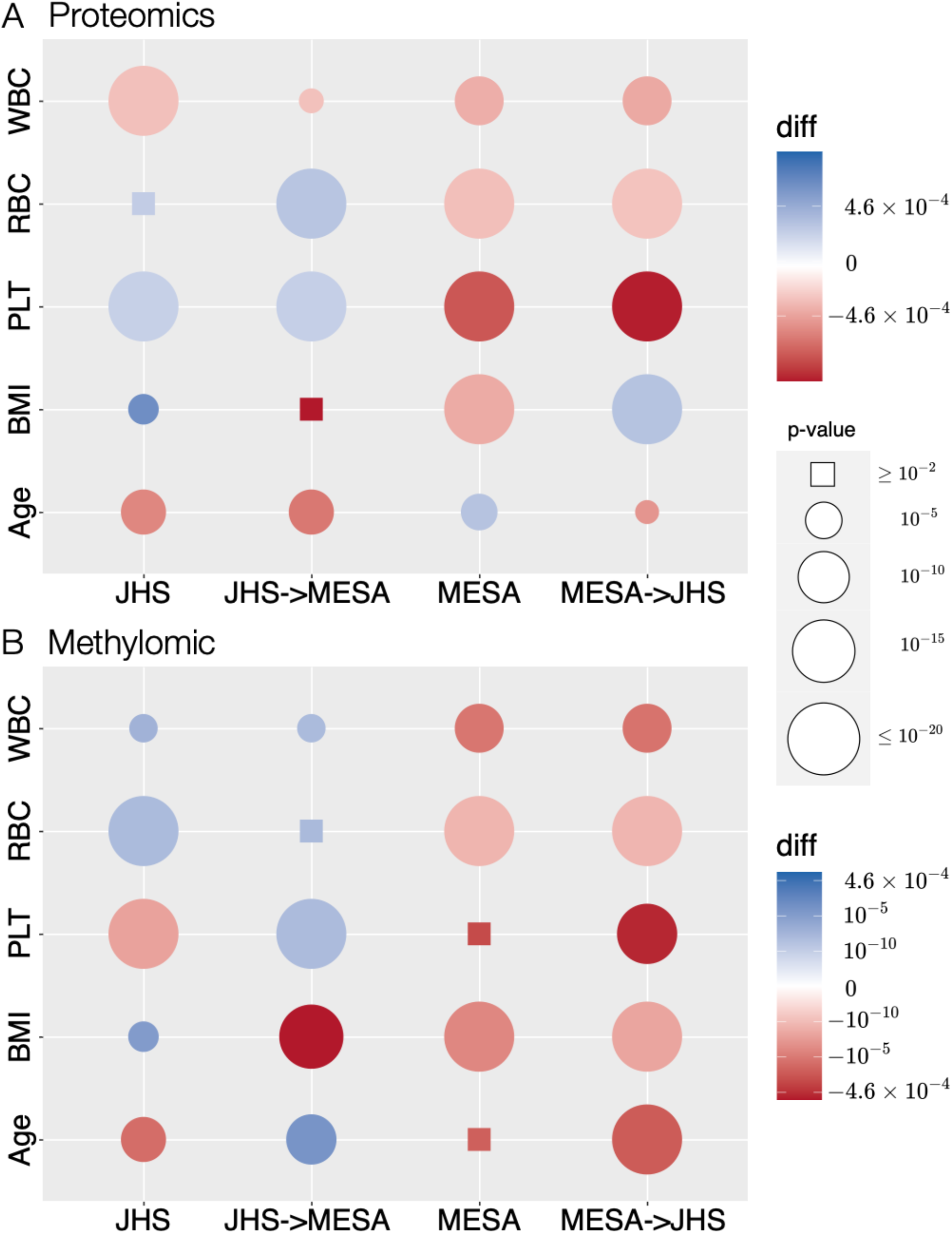
Comparison of SSCCA and SSMCCA. **(A)** proteomics, and **(B)** methylomics. Each row corresponds to a phenotype (from bottom to top, Age, BMI, WBC, RBC, and PLT). Circle size reflects the significance of the difference between two methods. Color reflects the difference between the variation of phenotype explained by SSCCA and our SSMCCA. Therefore, a larger circle means that the difference between two methods is more significant. Note that we use rectangles for insignificant difference with p > 0.01. Red means that our SSMCCA explains more phenotypic variation while blue means that SSCCA explains more.

#### Biologically Meaningful Features Detected by SSMCCA

We applied SSMCCA to three assays – proteomics, methylomics, and genotypes – from MESA to obtain 50 CVs for each assay, and then used standard regression models to assess associations with phenotypes – age, BMI, WBC, RBC, and PLT. CV-phenotype pairs were considered to be significantly associated when p-value < 1E-4 (Bonferroni correction), adjusting for covariates detailed in **S2 Table**. In MESA, we identified 58 significant CV-phenotype pairs, and 5 of them were validated in JHS with the same p-value threshold of 8.62E-4 and same direction of effect (**S3 Table**). For example WBC and proteomics CV3 were strongly associated in both cohorts (p-value = 2.7E-15 in MESA, 6.8E-16 in JHS, **S3 Table**). Features with high absolute weight coefficients in this CV (**S5 Table**) are biologically relevant for WBC. For example, stem cell factor soluble receptor, which has the highest weight, is known to play a key role in hematopoiesis [25]. Lipocalin 2, with the second highest weight, has been reported to be associated with human neutrophil granules [26].

For each phenotype, we then assembled all features from each assay with non-zero weight for phenotype-associated CVs in **S3 Table**, and annotated each feature to a gene, on which we performed pathway enrichment analysis (described in methods section). For comparison, we also performed the same pathway enrichment analysis using features individually associated with each phenotype, where association is declared when FDR < 0.05 for each assay-phenotype-cohort combination. Comparing these two sets of pathway enrichment results, we found several pathways only revealed (p.adjust < 0.05) by our SSMCCA, including the growth factor binding GO term, DisGeNET progressive chronic graft-versus-host disease (GVHD) and polypoidal choroidal vasculopathy genesets. All of these pathways have been reported to be related to BMI in previous literature [27–30].

## Discussion

Large quantities of data across multiple omics (transcriptomics, proteomics, metabolomics, genomics, methylomics, etc) modalities is currently being generated, for example through efforts funded by NIH’s Precision Medicine Initiative [24,31] as well as other large federally funded studies [32]. These high dimensional and complex multi-assay data are unfortunately still too often analyzed only separately (e.g., applying PCA separately to genotype, gene expression, or methylation data) or in a pairwise manner (for example mQTL analysis examining relationships between genome and methylome, or pQTL analysis examining relationships between genome and proteome). Many innovative methods have been proposed (https://github.com/mikelove/awesome-multi-omics [accessed on *2022-07-25*]) for integrative analysis but evaluations in large-scale real omics data are still lacking, with fewer impartial appraisals available to guide method choice in practice.

In the work presented here, we apply CCA-based methods to complex multi-omics datasets to assess their capabilities and limitations. In particular, for the widely used PMA implementation of the SMCCA methods, we identified two limitations: non-orthogonal CVs and inability to accommodate more than two assays for supervised analysis. We provide method extensions, SMCCA-GS and SSMCCA, to address the two limitations. Applying SMCCA-GS to real data in MESA and JHS, we found that CVs are consistent and transferable across cohorts, suggesting that CVs capture constitutive biological relationships shared across cohorts, and are not driven primarily by assay-specific technical variation. This cross-cohort consistency, to our knowledge, has not been well explored in the literature and has important implications for making method choices (e.g., CCA vs PCA) for multi-omics data with or without extensive assay-specific batch effects.

Importantly, our CCA-based analyses reveal that blood cell indices are substantially associated with multiple omics assays including methylomics and proteomics. The former association has been widely appreciated and exerted paradigm-shifting impact on analysis: in methylation association studies,white blood cell composition is adjusted for in methylation analyses in standard practice. The latter association, where CVs from proteomics data showed even more pronounced association with blood cell indices, has been under-appreciated [21,33–35]. Our findings indicate that blood cell composition should be accounted for in protein association studies, similar to what is standard practice for methylation studies.

As demonstrated in **Fig 5**, our SMCCA-GS is in some cases more useful than PCA in explaining variability in phenotypes, using an identical number of PCs/CVs. However, there are also many cases where the methods are nearly equivalent. We hypothesize that our SMCCA-GS demonstrates more consistent advantages in explaining trait variability in JHS versus MESA due to the presence of more substantial JHS batch effects. Due to funding limitations, JHS proteomics and metabolomics data was generated in multiple batches across several years, while the MESA data used here was generated concurrently, funded by NHLBI’s TOPMed program. Thus, for proteomics in particular, more batch effects are anticipated in JHS; our SMCCA-GS is particularly advantageous in cases where there is increased assay-specific technical variation.

We note that CCA-based methods as implemented in our analyses still have several key limitations. Notably, we had to considerably reduce the dimensionality of methylation array and sequencing data in order for our CCA-based method to be computationally feasible. We thus are not capturing anywhere close to the maximal information potential from these assays. Further innovation can make our SMCCA-GS more computationally feasible for large-scale datasets. Recently developed methods allow for efficient calculation of generalized CCA solutions across reduced dimensions of each distinct assay, which alleviates some of the computational issues that arise, though sparse identification of individual omics features from the original assay data may still be desired [36].

## Methods

### Cohorts

All participants included in this analysis provided written, informed consent for use of genetic and multi-omics data, and all study protocols conform to the 1975 Declaration of Helsinki guidelines. The JHS and MESA studies were approved by the Institutional Review Boards of all participating institutions.

### JHS

JHS recruited 5,306 African American participants from the Jackson, Mississippi, metropolitan tri-county area (Hinds, Madison, and Rankin) into a prospective, community-based cohort designed to investigate risk factors for cardiovascular disease among African Americans [37–39]. Demographics of JHS individuals involved in the analysis is in **S1 Table A**.

Multi-omics data utilized in JHS analyses including methylomics (n = 1,750, Illumina MethylationEPIC BeadChip array) [40] and proteomics (n = 2,144, SOMAscan 1.3k array) [21], both from the baseline visit, and whole genome sequencing (WGS) data as described below. Traits examined include age, sex, BMI, and hematological traits (WBC, RBC, and PLT). We limited our analyses in JHS to individuals with complete data for proteomics, methylomics, and traits examined (total n = 881, **S1 Fig A**).

### MESA

The MESA study was initiated in July 2000 to investigate the prevalence, correlates, and progression of subclinical cardiovascular disease (CVD) in a population-based sample of 6,814 men and women aged 45–84 years. The cohort was selected from six US field centers. Based on self-reported race/ethnicity, approximately 38% of the cohort are White, 28% African-American, 23% Hispanic, and 11% Chinese American. More demographic information of MESA individuals involved in the analysis is in **S1 Table B**.

Longitudinal multi-omics data was generated in MESA through a pilot program from NHLBI’s Trans-Omics for Precision Medicine Initiative (TOPMed) at exam 1 (2000-2002) and exam 5 (2010-2011), including ~ 1,000 participants for each exam with methylomics data (Illumina MethylationEPIC BeadChip array) [41] and proteomics (SOMAscan 1.3k array) [21,22]. WGS data are described below. Basic covariates examined include age, sex, BMI, recruitment site, self-reported race/ethnicity, and the same hematological traits as in JHS. We limited our analyses in MESA to individuals with complete data for proteomics, methylomics, and phenotypes examined (total n = 777, **S1 Fig B**). Use of the same platforms for multi-omics assessment as in JHS allowed comparison analyses for CVs derived by SMCCA-GS or SSMCCA across cohorts.

### Whole Genome Sequencing (WGS) Data

Genotypes are derived from TOPMed WGS data (freeze 8). Data harmonization, variant discovery, and genotype calling were previously described [24,42]. In our analysis, to reduce data dimensionality, SNPs associated with blood cell traits from Chen et al. (2020) [43] were extracted, and highly correlated (linkage disequilibrium *r*^2^ > 0.8 where *r*^2^ is estimated by the squared Pearson correlation between genotype vectors) variants within JHS or MESA were removed. Population principal components calculated by PC-AiR [44] were adjusted for as covariates. In addition, for WBC, we additionally adjusted for the Duffy null polymorphism (SNP rs2814778 at chromosome 1q23.2) [45].

### Initial quality control (QC) and transformation of multi-omics data

In both cohorts, we applied QC on each assay including sample outlier removal and feature filtering. For each protein in the proteomics data, we first applied log transformation, followed by inverse normal transformation. After QC, we had 1,317 proteins measured in both cohorts, which made validation across cohorts straightforward.

Methylomics of JHS [40] was normalized using the “noob” normalization method implemented in minfi R package [58,59]. We further removed batch, plate, row, and column effects using the ChAMP R package [46]. For MESA methylation data, we excluded samples with (1) call rate < 95%; (2) sex mismatches; and (3) concordance between SNP probes and genotypes < 0.8. Methylation levels were marked as missing when the detection p-value was > 0.01, and we imputed these missing values using ChAMP R package [46], as our CCA-like methods cannot accommodate missing data. For both JHS and MESA, CpG sites whose probes overlap any SNP with minor allele frequency (MAF) > 1% were also excluded [47]. After QC, we had 754,767 and 741,727 CpG sites for MESA and JHS respectively. For building validation across cohorts, we only kept the 721,334 CpG sites which passed QC in both cohorts.

Finally, we only kept samples with complete data including proteomics, methylomics, and phenotypes (age, BMI, WBC, RBC, PLT, site, race, sex for MESA; age, BMI, WBC, RBC, PLT, sex for JHS), which led to 881 samples for JHS and 777 samples for MESA. We further identified sample outliers by PCA-IQR plot (**S2 Fig**). Four outliers in JHS – one sample with the largest proteomics IQR (wedge pointed on **S2 Fig A**) and three samples with largest methylomics IQR (wedges pointed on **S2 Fig B**) – were removed; and three outliers in MESA were removed – all three with largest methylomics IQR (wedges pointed on **S2 Fig D**).

For each assay, we removed the sex chromosome related proteins and CpG sites. We further removed features that are highly correlated [48], at a squared Pearson correlation 0.8 threshold. For methylomics, we calculated Pearson correlation using the Python package Deep Graph [48] and after removing highly correlated, further kept 10k CpG sites with the highest variance for the computational efficiency. Our CCA-based methods are computationally intensive. For example, even with these 10k CpG sites (~1.3% of all available CpG sites), on a single core of E5-2680 v3 @ 2.50GHz, the wall time of calculating 50 CVs with our SMCCA-GS on proteomics and methylomics is about 8 ~ 14 hours for MESA (774 samples) and about 8 ~ 20 hours for JHS (877 samples); with 20k CpG, the wall time is about 14 ~ 36 hours for MESA and 16 ~ 47 hours for JHS. For validating our variance-based feature selection strategy, we also performed the same analysis as **Fig 4, 5** on proteomics and all ~700k CpG sites. The results (**S3 & S4 Fig**) show similar patterns as those from top 10k CpG sites (**Fig 4, 5**).

### Modified Gram-Schmidt Strategy

With PMA implementation, we observed that with our real data where features have complex correlation structure, the weight vectors are sometimes correlated. To mitigate this non-orthogonality issue, we adopt a strategy inspired by Woojoo et al., (2011) [15]. In our implementation, we infer CVs sequentially and remove the effects of the former CVs from the input matrix before calculating weights for the next CVs. In particular, we first follow the PMA approach to generate weights for CV1’s of all assays, update input matrices following Eq (2) as the new inputs for calculating weights for CV2’s, and sequentially update until we obtain pre-specified numbers of CVs. Eq (1) and Eq (2) show the procedures for inferring the (*j*+1)’s CVs with input matrices {*X_ij_*}_*i*=1,…,*S*_.

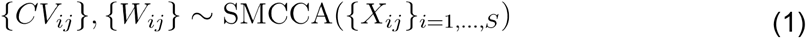

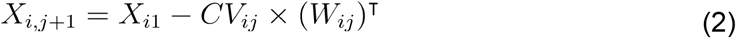

Specifically, {*X*_*i*1_}_*i*=1,…,*S*_ are original input matrices for assay *i*= 1,…, S where S is the total number of assays, from which *W*_*i*1_’s and *CV*_*i*1_’s, the first weights and CV1’s, are inferred by SMCCA implemented in the PMA R package.

### Pathway Enrichment Analysis (add more info about the functions and settings)

For each CCA-prioritized feature of each assay, we first mapped them to genes, and then performed pathway enrichment analysis on these genes utilizing three databases – DisGeNET [22–24] (enrichDGN function in DOSE R package, with default settings), Gene Ontology (GO) [25,26] (enrichGO function in clusterProfiler R package, with default settings) and Kyoto Encyclopedia of Genes and Genomes (KEGG) [27–29] (enrichKEGG function in clusterProfiler R package, with default settings). For methylomics, we mapped CpG sites to the nearest genes using annotations released by Illumina. For proteomics, we mapped proteins to genes using annotations released by SomaScan. For background genes in the enrichment analysis, we included genes annotated from features that are associated with outcome, identified in the feature selection step of our SSMCCA (detailed in Section “Extending Supervised Sparse CCA to Supervised Sparse Multiple CCA” above).

## Code Availibility

Our modified SMCCA-GS and SSMCCA functions are available at https://github.com/zjgbz/SMCCA-GS_SSMCCA.

## Author Contributions

**Conceptualization:** Yun Li, Michael I. Love, Laura M. Raffield.

**Data curation:** Min-Zhi Jiang, Laura M. Raffield

**Formal analysis:** Min-Zhi Jiang.

**Funding acquisition:** Yun Li, Michael I. Love, Laura M. Raffield.

**Investigation:** Yun Li, Michael I. Love, Laura M. Raffield, Min-Zhi Jiang.

**Methodology:** Yun Li, Michael I. Love, Laura M. Raffield, Min-Zhi Jiang.

**Project administration:** Yun Li, Michael I. Love, Laura M. Raffield.

**Resources:** Yun Li, Michael I. Love, Laura M. Raffield, James G. Wilson, Stephen S. Rich, Jerome I. Rotter.

**Software:** Min-Zhi Jiang.

**Supervision:** Yun Li, Michael I. Love, Laura M. Raffield.

**Visualization:** Min-Zhi Jiang, Jiawen Chen.

**Writing – original draft:**Yun Li, Michael I. Love, Laura M. Raffield, Min-Zhi Jiang.

**Writing – review & editing:** Min-Zhi Jiang, François Aguet, Kristin Ardlie, Jiawen Chen, Elaine Cornell, Dan Cruz, Peter Durda, Stacey B. Gabriel, Robert E. Gerszten, Xiuqing Guo, Craig W. Johnson, Silva Kasela, Leslie A. Lange, Tuuli Lappalainen, Yongmei Liu, Alex P. Reiner, Josh Smith, Tamar Sofer, Kent D. Taylor, Russell P. Tracy, David J. VanDenBerg, James G. Wilson, Stephen S. Rich, Jerome I. Rotter, Michael I. Love, Laura M. Raffield, Yun Li.

---

**S1 Fig.**
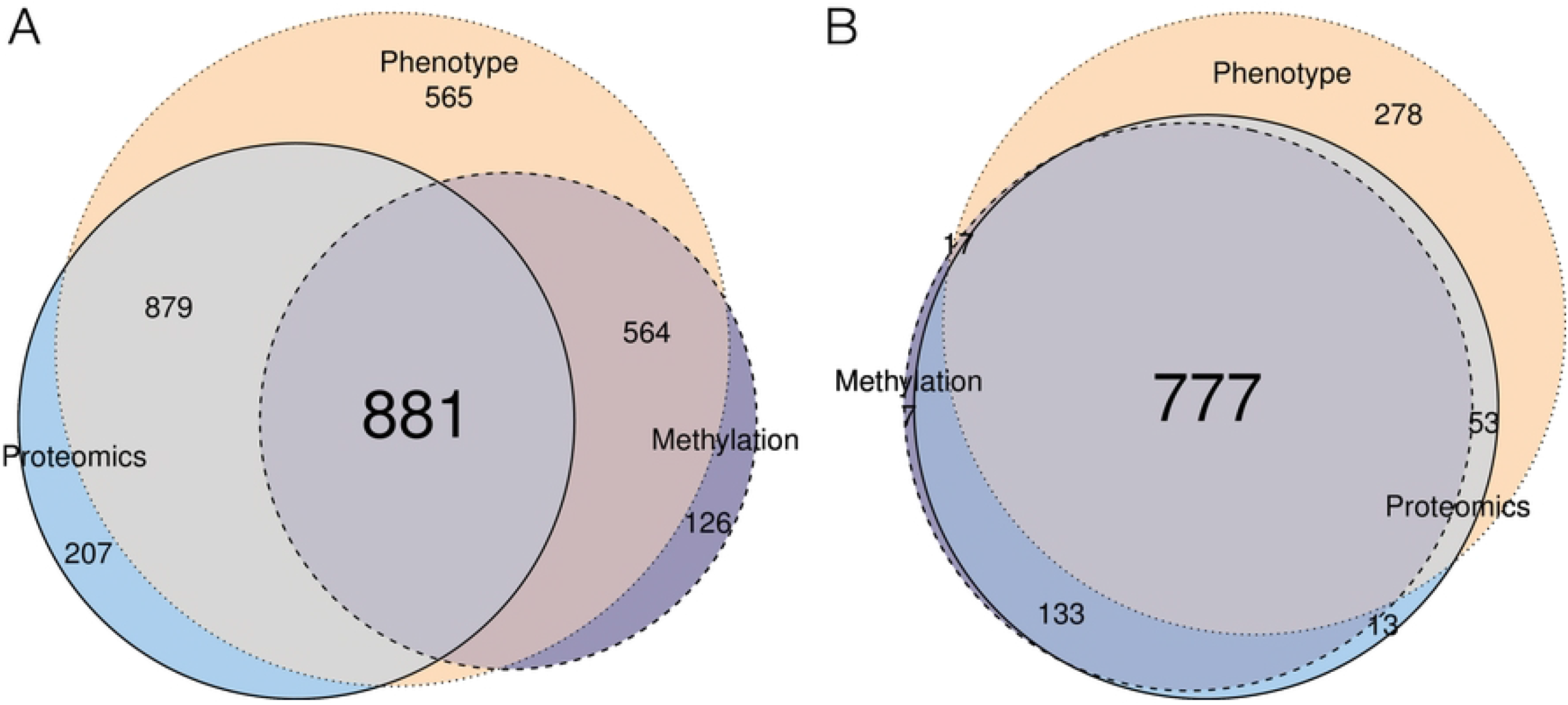

**S2Fig.**
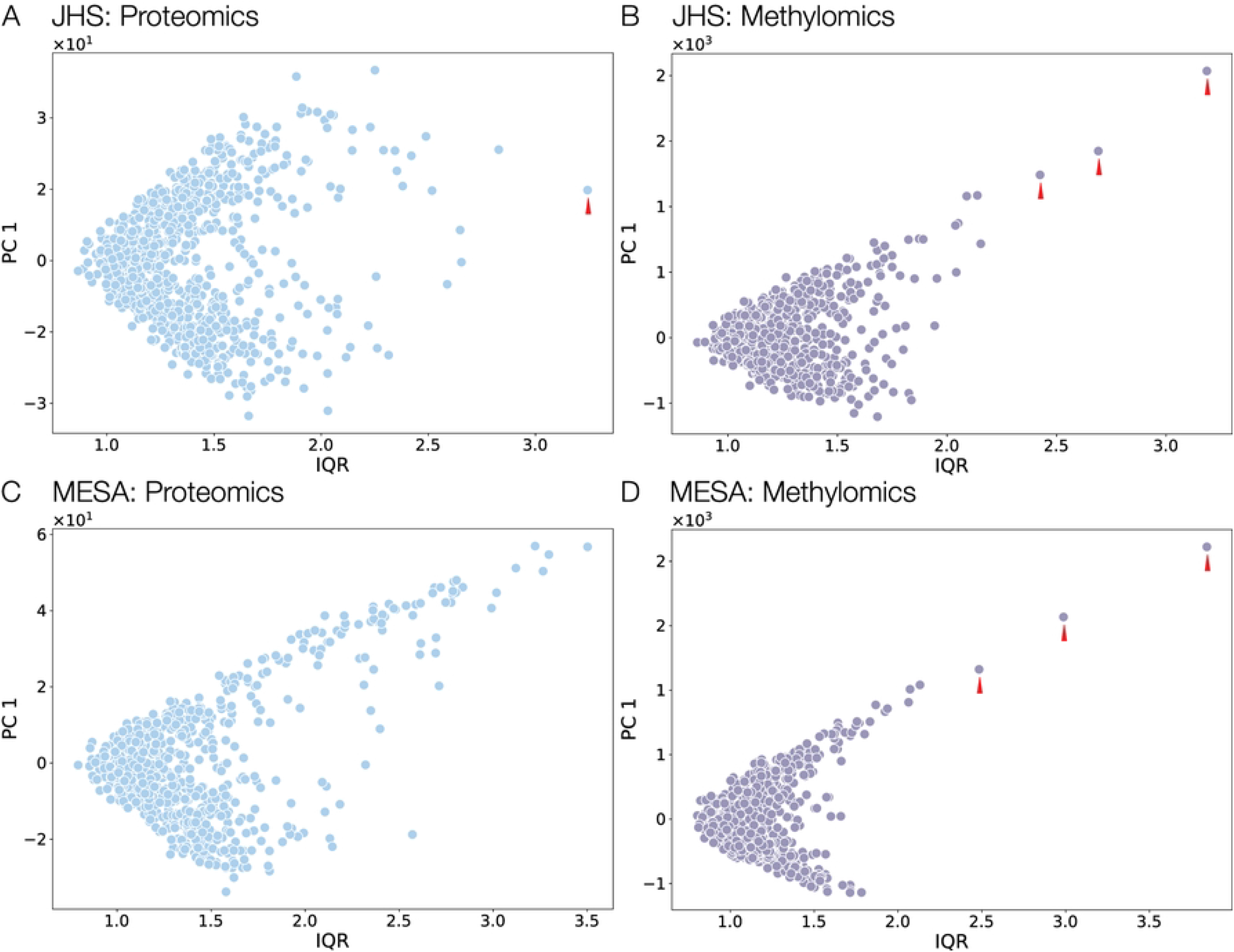

**S3Fig.**
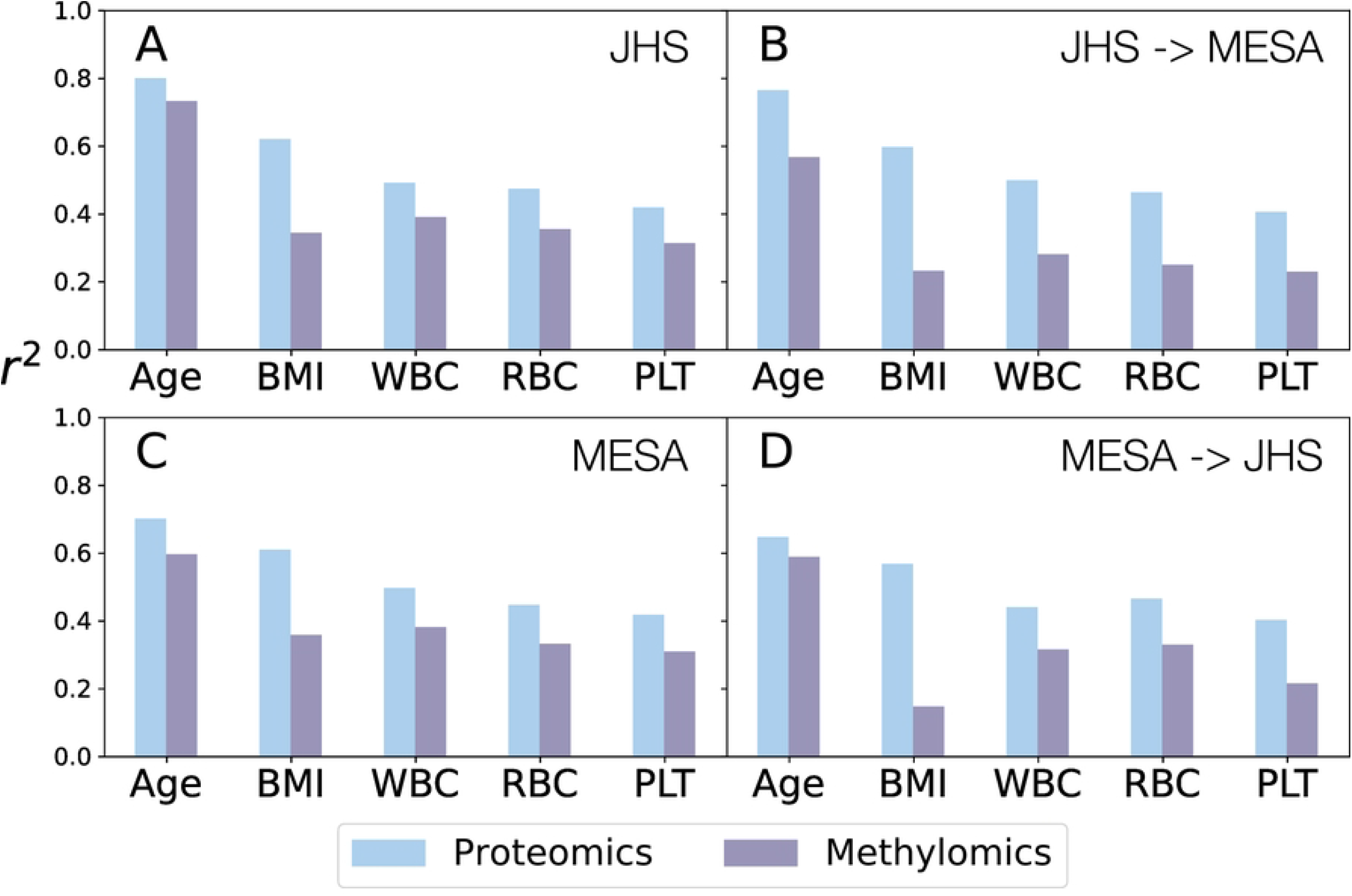

**S4Fig.**
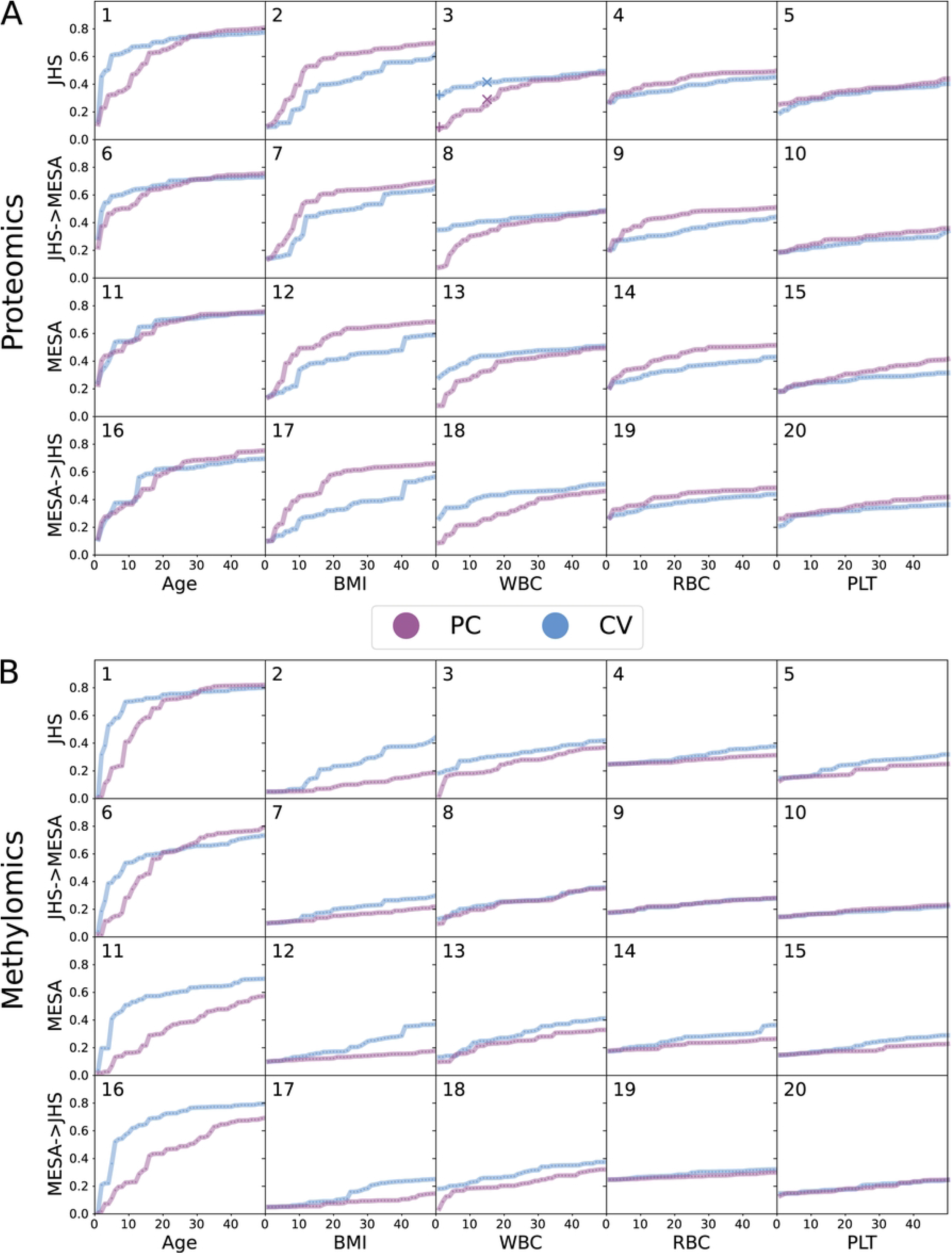

